# Genomic analysis of BCG unresponsive non-muscle-invasive bladder cancer identifies drivers of sensitivity to intravesical Gemcitabine/Docetaxel

**DOI:** 10.64898/2026.05.10.724123

**Authors:** Kendrick Yim, Matias Vergara, Jihyun Lee, Brendan Reardon, Jihye Park, Kevin Melnick, Timothy N Clinton, Matthew Mossanen, Graeme S Steele, Jessica Bolduc, Michelle Hirsch, Natalie Rizzo, Chin-Lee Wu, Matthew Wszolek, Keyan Salari, Adam Feldman, Adam S Kibel, Kent W Mouw, Eliezer Van Allen, Mark A Preston, Filipe LF Carvalho

**Author notes:** <u>Corresponding Authors:</u> Filipe LF Carvalho, MD, PhD, Brigham and Women’s Hospital, Department of Urology, 45 Francis Street, ASB II, 3^rd^ Floor. Boston, MA, 02115, Mark Preston, MD, MPH, Brigham and Women’s Hospital, Department of Urology, 45 Francis Street, ASB II, 3^rd^ Floor. Boston, MA, 02115. Contributed equally (Kendrick Yim, Matias Vergara, Filipe LF Carvalho).

## Abstract

**Background and Objectives:** Intravesical gemcitabine/docetaxel (Gem/Doce) is an effective therapy for Bacillus Calmette– Guérin (BCG)-unresponsive non–muscle-invasive bladder cancer (NMIBC), achieving 50% complete responses at 2 years. However, the genomic determinants underlying response and resistance to Gem/Doce remain poorly defined. Our objective was to define the mutational landscape of BCG-unresponsive NMIBC and nominate genomic features associated with response or resistance Gem/Doce.

**Methods:** Patients with BCG-unresponsive NMIBC treated with Gem/Doce were classified as responders (recurrence-free survival [RFS] >12 months) or non-responders (RFS <12 months). Whole-exome sequencing was performed on tumors prior to Gem/Doce treatment (n=23). Single nucleotide variants were identified and annotated using a Cancer Genome Analysis pipeline. Copy number alterations were inferred with ABSOLUTE, and clonal architecture was reconstructed using PhylogicNDT.

**Key Findings and Limitations:** Responders demonstrated significantly prolonged time to high-grade recurrence (3.5 vs 42 months, p<0.001) and cystectomy compared with non-responders (9.5 months vs not reached; p<0.001). Non-responders exhibited higher tumor mutational burden (13.66 vs 8.71; p=0.02) and more frequent whole-genome doubling (2/2 non-responders vs 0/1 responders; p=0.33). Phylogenetic analyses revealed clonal *BAP1* and subclonal *BRCA2* mutations in responders, whereas non-responders harbored clonal *FGFR3* mutations. Limitations include small sample size and retrospective design.

**Conclusions and Clinical Implications:** Distinct genomic features underlie differential response to Gem/Doce in BCG-unresponsive NMIBC. In responders, alterations in DNA repair pathways (e.g., *BRCA2*) may sensitize tumors to chemotherapy, while non-responders with FGFR3 mutations may benefit from alternative targeted strategies. These findings warrant validation in larger cohorts and support the development of biomarker-driven clinical trials.

**Patient summary:** In this report we analyzed bladder tumors and found that some tumors respond well to treatment because they have defects in repairing DNA, making them more vulnerable to chemotherapy. In contrast, tumors that do not respond to chemotherapy harbor different genetic changes that help them survive and grow. These findings may help physicians choose more effective and personalized treatments in the future.

## Introduction

Approximately 85,000 people will be diagnosed with bladder cancer in the United States in 2026 and 75% of cases will be non-muscle invasive tumors[1]. Transurethral resection followed by intravesical Bacillus Calmette-Guerin (BCG) remains the standard of care for high-risk non-muscle invasive bladder cancer (NMIBC); however, 25% of patients do not respond to BCG and 50% will recur within 5 years[2]. Because radical cystectomy with urinary diversion is associated with high morbidity and complications, there has been tremendous interest in developing bladder-sparing intravesical treatments for patients with BCG-unresponsive NMIBC.

Sequential intravesical gemcitabine and docetaxel (Gem/Doce) has demonstrated promising efficacy in the treatment of BCG-unresponsive NMIBC with 2-year recurrence-free survival rates of 50% and lower risk of progression to MIBC when compared to salvage BCG[3-5]. Gemcitabine is a nucleoside analog that inhibits DNA synthesis, and docetaxel is a microtubule stabilizing agent that disrupts cell division. Together, these agents act synergistically to induce cytotoxicity and mitotic errors in bladder cancer cells[6, 7]. Recent studies suggest that Gem/Doce has a favorable oncologic control and tolerability in comparison to newer and more costly FDA-approved therapies such as Pembrolizumab (12-month disease free rate of 18%), nadofaragene firadenovec (12-month complete response rate of 17%), and nogapendekin alfa inbakicept + BCG (12-month complete response rate of 45%)[8-10]. However, there is no established algorithm for sequencing of these treatments after BCG failure and no validated predictive or prognostic biomarkers to guide treatment-decision making. Consequently, patients may be exposed to ineffective therapies and associated toxicities, experience delays in more effective treatments, ultimately leading to a higher risk of progression to muscle-invasive disease.

Recent efforts have been focused on comprehensive genomic characterization of NMIBC to better understand genomic drivers of disease biology, including progression to MIBC[11, 12] and response to BCG[13, 14]. However, no studies to date have examined genomic features or nominated predictive biomarkers for response to Gem/Doce. Here, we analyzed tumor whole exomes from a cohort of patients with BCG-unresponsive NMIBC treated with intravesical Gem/Doce in order to nominate genomic alterations that sensitize BCG-unresponsive NMIBC to Gem/Doce and provide a biological rationale for sequential intravesical treatments.

## Materials and Methods

### Patients and samples

With Institutional Review Board approval (2021P002957/GC24-02), we reviewed records from patients with NMIBC who received sequential intravesical Gem/Doce after disease recurrence following at least one induction course of BCG (≥5 instillations) and ≥2 re-induction or maintenance BCG instillations at Brigham and Women’s Hospital. Details regarding patient selection, Gem/Doce sequential therapy, and surveillance protocols have been previously described[4]. For each case, a specimen from a transurethral bladder tumor resection (TURBT) performed after BCG failure and prior to Gem/Doce was reviewed by a board-certified genitourinary pathologist (M.H. and C.L.W). Cores from FFPE blocks with areas with high tumor density were collected for DNA isolation (n= 23). Peripheral blood was used to isolate germline genomic DNA in three patients for matched tumor-normal DNA pairs. Demographic and clinical characteristics were collected and integrated with genomic findings. Patients were defined as “responders” to Gem/Doce when HG-RFS was ≥ 12 months after first Gem/Doce instillation or “non-responders” when HG recurrence (HG-RFS <12 months) or progression occurred within 12 months after first Gem/Doce instillation[15].

### DNA extraction and WES

DNA was isolated from FFPE punch cores from TURBT specimens while matched germline DNA was purified from peripheral blood. Samples with at least 100 ng of dsDNA were used to generate libraries for whole-exome sequencing (WES). The sequencing protocol has been described previously[16].

### Data processing and analysis

A WES characterization pipeline developed at the Broad Institute was used to identify, filter, and annotate somatic mutations[16]. BAM files from 20 samples without matched germline DNA were processed with the Cancer Genome Computational Analysis (CGA) tumor-only pipeline, and samples with matched tumor/germline DNA pairs were processed with the original CGA pipeline. We also implemented a variant filtering strategy to remove common germline variants by filtering those that appear in at least 10 alleles across any subpopulation within the ExAC database, now a part of gnomAD[17]. Cross-individual contamination was estimated using ContEst[18], and only samples with <5% estimated contamination were included in the study. Single nucleotide variants (SNVs) in targeted exons for each sample were identified using MuTect[19], and indels identified using MuTect2[20] and Strelka[21]. Genomic alterations were then annotated using Oncotator[22]. Possible sequencing artifacts were filtered using a pipeline developed at the Broad Institute[23]. Alterations in prespecified frequently mutated NMIBC genes and mutations in DNA damage repair genes were identified[20]. Gene mutations across the cohort were summarized in co-mutation plots[24].

### Total and allelic copy numbers

Allele-specific copy number at a gene level were inferred using ABSOLUTE on tumor/normal pairs for allele-specific copy number calls[25].

### Mutation signature analysis

Somatic SNVs were used to infer mutational signature activities using SigProfilerAssignment[26] and signature decomposition was performed against the COSMIC v3.4 reference signature set[27]. Differences in SNVs between groups were calculated using Wilcoxon rank sum test and permutation Welch test.

### Phylogenic analysis

The clonal architecture of tumors with matched tumor-normal pairs was reconstructed using PhylogicNDT[28]. Cluster module estimates the number of distinct tumor subclones and assigns individual mutations to their respective subpopulations

### Data availability and code

Code will be publicly available on GitHub (https://github.com/CarvalhoFilipeL) at the time of publication. To be publicly available, called variants are being deposited in the cBioPortal for Cancer Genomics (https://www.cbioportal.org/).

## Results

### Clinical Characteristics of BCG unresponsive patients

We identified a total of 23 patients with recurrent NMIBC after BCG who were subsequently treated with intravesical Gem/Doce. BCG-unresponsive disease was defined as ≥ 5 of 6 induction doses of BCG and ≥2 doses of maintenance or re-induction BCG. Clinical characteristics of the cohort are summarized in Table 1. Thirteen patients were designated as “responders” to Gem/Doce (HG-RFS >12 months), while 10 were designated as “non-responders” to Gem/Doce (HG-RFS <12 months). The median number of prior BCG instillations in the responder and non-responder groups was 12 and 13.5, respectively (Wilcoxon rank-sum test p= 0.97). Pre-treatment tumor stage was similar between responders and non-responders: in responders 15% CIS, 54% Ta-HG, and 31% T1HG; and in non-responders 10% CIS, 10% Ta-LG, 80% Ta-HG, and no T1HG (Chi-square test, p=0.18). Median time to HG-recurrence was 3.5 months in non-responders versus 42 months in responders (log rank p<0.0001, Fig. 1A). Median time to cystectomy was 9.5 months in non-responders and not reached in responders (log rank p=0.0006 Fig. 1B).

**Table 1.** Demographic, pathologic, and treatment features of patients undergoing gemcitabine/docetaxel intravesical therapy.

|  | Responders (n=13) | Non-responders (n=10) |
| --- | --- | --- |
| Median Age, years (IQR) | 70 (67- 75) | 73 (69 – 76) |
| Median follow up, months (IQR) | 40 (28 – 44) | 44 (30 – 58) |
| Gender (%) |  |  |
| Male | 8(62%) | 7 (70%) |
| Female | 5 (38%) | 3 (30%) |
| Race (%) |  |  |
| White | 12 (92%) | 9(90%) |
| Hispanic | 1 (8%) | 1 (10%) |
| Smoker (%) |  |  |
| Yes (current or former) | 7 (54%) | 6 (60%) |
| Never | 6 (46%) | 4 (40%) |
| Median number prior BCG instillations (IQR) | 12 (6 – 20) | 13.5 (7 – 17.5) |
| Pre-Treatment Stage (%) |  |  |
| CIS alone | 2 (15%) | 1 (10%) |
| Ta-LG | 0 (0%) | 1 (10%) |
| Concurrent CIS | 0 (0%) | 0 (0%) |
| Ta-HG | 7 (54%) | 8 (80%) |
| Concurrent CIS | 0 (0%) | 2 (20%) |
| T1-HG | 4 (31%) | 0 (0%) |
| Concurrent CIS | 0 (0%) | 0 (0%) |
| Median number of gem/doce instillations (IQR) | 18 (13 – 22) | 6 (6 – 6) |
| Post-Treatment Stage (%) |  |  |
| No recurrence | 9 (69%) | 0 (0%) |
| CIS alone | 2 (15%) | 2 (20%) |
| Ta-HG | 1 (8%) | 6 (60%) |
| Concurrent CIS | 0 (0%) | 2 (20%) |
| T1-HG | 1 (8%) | 2 (20%) |
| Concurrent CIS | 0 (0%) | 1(10%) |
| Median Time to HG recurrence, months (IQR) | 42 (32 – NA) | 3.5 (2 – NA) |
| Received cystectomy, n (%) | 2 (16%) | 7 (70%) |
| Median time to cystectomy, months (IQR) | NA (NA – NA) | 9.5 (5 – NA) |

**Figure 1.**
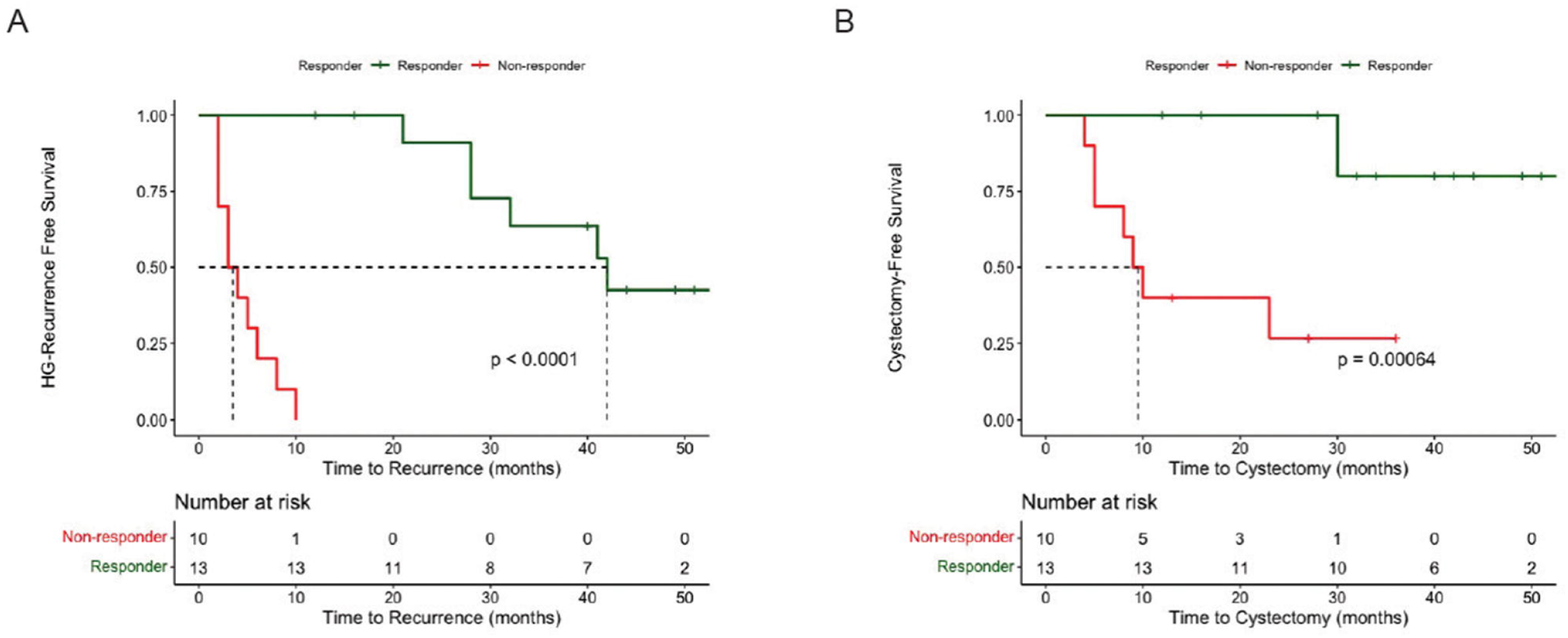
High-grade recurrence free survival (A) and cystectomy-free survival (B) in the BCG-unresponsive cohort in patients with extreme response or resistance (Non-responders) to gemcitabine/docetaxel. Survival times were calculated from the date of first dose of gemcitabine/docetaxel.

### Genomic landscape of responders versus non-responders to Gem/Doce

Given the difference in tumor response and clinical outcomes between Gem/Doce responders and non-responders, we performed whole-exome sequencing of BCG unresponsive tumors prior to treatment with Gem/Doce to nominate genomic correlates of recurrence. The cohort comprised 23 tumors, 3 matched tumor/normal pairs and 20 tumor-only samples. The mean target coverage was 172x for somatic tumor DNA for the 20 tumor-only samples. For the 3 matched tumor/normal pairs, mean target coverage was 349x for somatic tumor DNA and 210x for germline DNA. For the tumor-only samples, a total of 10,719 variants were identified, including 6,232 missense, 453 nonsense, 208 frameshift, and 2,822 synonymous variants. For the tumor-normal samples, a total of 1,645 variants were identified, including 597 missense, 40 nonsense, 13 frameshift and 230 silent mutations (Supplementary Table 1).

We first evaluated genes and copy number alterations known to be recurrently mutated in NMIBC[13, 29]. We identified mutations in several genes implicated in bladder cancer in both responders and non-responders (Fig. 2A), including *TP53* (11/23, 48% tumors), *KDM6A* (10/23, 43%tumors), *FGFR3* (6/23, 26%tumors), *KMT2A* (4/23, 17% tumors), *KMT2D* (8/23, 35% tumors), and *ARID1A* (5/23, 22% tumors). In patients with paired germline and tumor DNA, copy number analysis revealed *CDKN2A* copy number loss in both responders and non-responders, but amplifications in *E2F3, FGFR3* and *MDM2* only present in non-responders. Moreover, whole-genome doubling was also only observed in non-responders (Fig. 2B, Suppl. Fig. 1).

**Figure 2.**
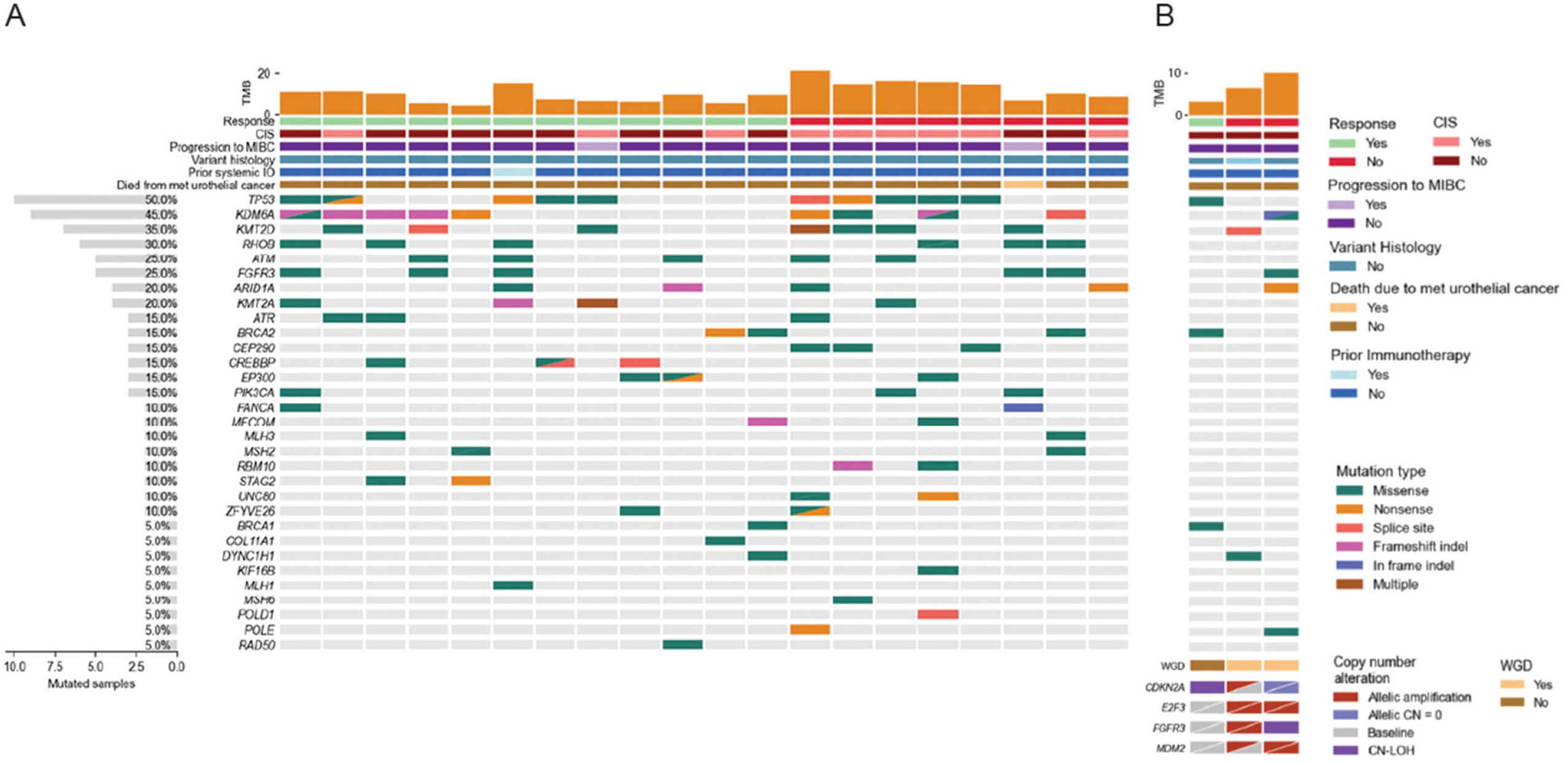
Mutational landscape of BCG-unresponsive cohort. Tumor mutational burden (TMB), response to gemcitabine/docetaxel, clinical characteristics, and alterations in select bladder cancer genes for each tumor in the cohort (n=23). Patients with tumor-only (A) and matched tumor/germline DNA (B) samples. Copy number alterations are represented for cases with matched tumor/germline pairs (bottom right). CIS = Carcinoma in situ. WGD = Whole genome doubling. CN = Copy number. LOH = Loss of heterozygosity.

Given that gemcitabine and docetaxel can lead to DNA damage in tumor cells, we assessed the mutational status of genes known to be involved in DNA damage and repair (DDR). Mutations in ATM, ATR, FANCA, and RAD50 were distributed across both groups without a clear enrichment in either group. Mutations in genes implicated in mismatch repair genes were also identified at low frequency and insufficient to elucidate statistically robust associations: *MLH1* (1/23 tumors); *MSH2* (2/23 tumors); and *MSH6* (1/23 tumors). However, mutations in *BRCA1* and *BRCA2* were exclusively present in responders (BRCA1 mutations in 2/23 tumors; BRCA2 mutations in 3/23 tumors; Fig. 2). Additionally, STAG2 and CREBBP mutations were also detected exclusively in responders (STAG2 mutations in 2/23 tumors; CREBBP mutations in 3/23 tumors). STAG2 is a core component of the cohesion complex and regulates homologous recombination repair, while CREBBP is a histone acetyltransferase involved in the recruitment of DNA repair factors[25,26]. Overall, non-responders to Gem/Doce had whole-genome doubling that promotes chromosomal instability and enables cancer cells to survive cytotoxic chemotherapy[27]. In contrast, responders harbor mutations in DDR genes suggesting that loss of DNA repair capacity may sensitize tumors to the DNA-damaging and mitotic stress-inducing effects of gemcitabine and docetaxel.

### Tumor mutational burden and mutational signatures

To further define the underlying mechanisms that distinguish responders from non-responders, we compared tumor mutational burden (TMB) and mutational signatures. We found that mean TMB was significantly higher in non-responders (13.66 mutations/Mb vs 8.71 mutations/Mb, Permutation Welch, p=0.02, Fig. 3A, Suppl. Fig. 2). Mutational signature analysis identified “clock-like” signatures (SBS1, SBS5) in all samples (Suppl. Fig. 3). We further explored APOBEC-related signatures (SBS 2 and SBS13) given their known role in NMIBC progression to MIBC [29-31]. We found that non-responders with APOBEC mutational signatures exhibited a higher TMB compared to responder tumors with APOBEC signatures (9.89 mutations/Mb vs 13.76 mutations/Mb, Permutation Welch, p=0.14, Fig. 3B). This association of the presence of an APOBEC signature and higher TMB in non-responders suggests a mechanistic link between these genomic features and a more aggressive clinical course of BCG-unresponsive NMIBC resistant to Gem/Doce.

**Figure 3.**
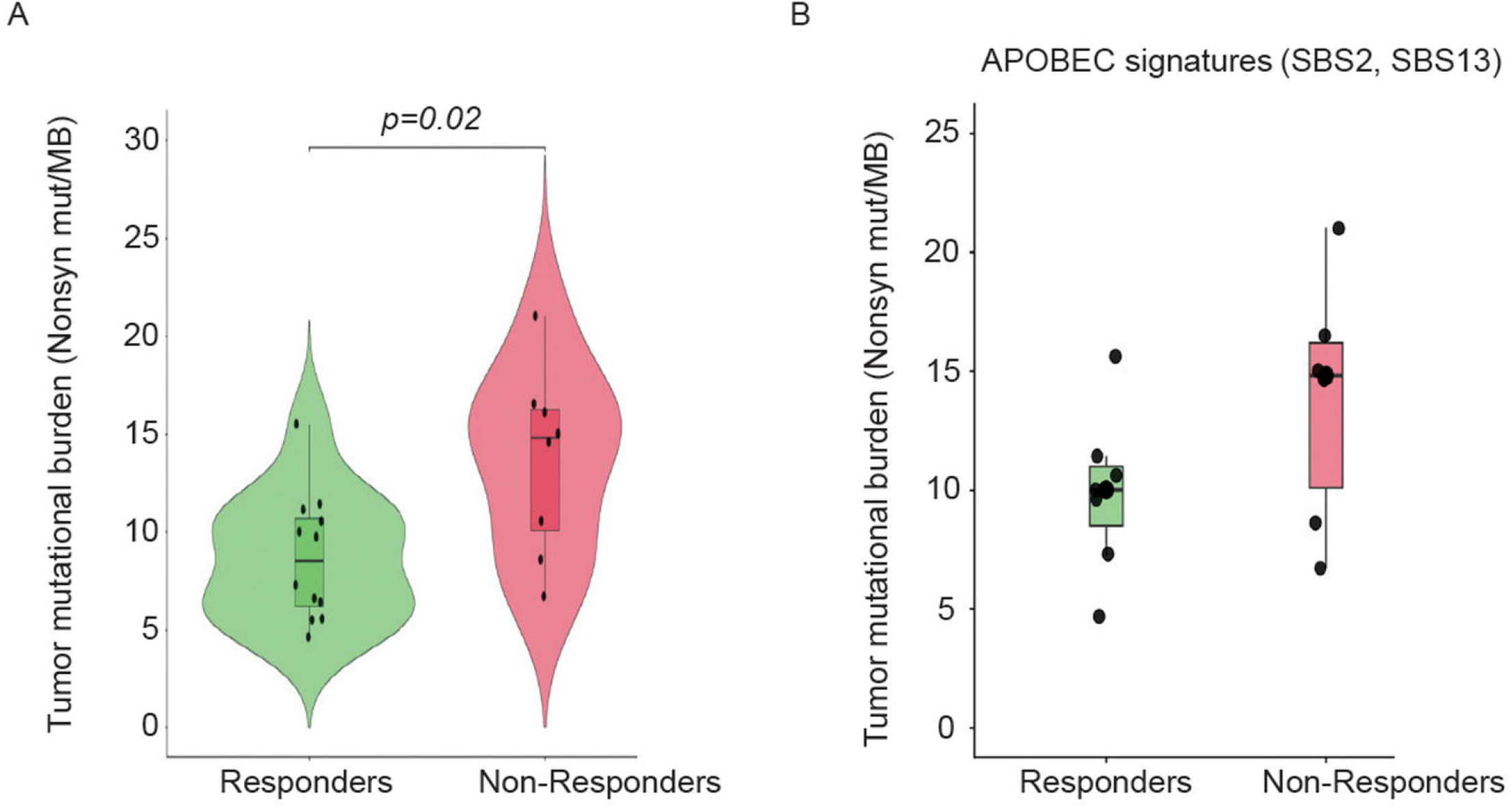
A. Tumor mutational burden (TMB) was significantly higher in non-responders compared with responders (Permutation Welch test, p= 0.02). B. APOBEC-mediated mutagenesis (mutational signatures SBS2 and SBS13) were identified in 7 responders and 7 non-responders. Non-responders with APOBEC signatures exhibited a higher TMB compared to responders with APOBEC signatures (mean 10.7 mutations/Mb vs 6.6 mutations/Mb, Permutation Welch test, p=0.14).

### Clonality and tumor evolution

Three patients (one responder, two non-responders) had paired tumor and normal exome sequencing which enabled clonal and genomic evolution analyses. Sample purity and allele-specific copy number data from ABSOLUTE was used to calculate the cancer cell fraction (CCF) of point mutations and construct a phylogenetic tree to dissect the clonal architecture and evolutionary trajectory of these tumors. We observed a clonal mutation (CCF = 1) in a driver cancer gene or tumor suppressor gene in all cases. In the responder case, the presence of a truncating *BAP1* p.W0R mutation as a clonal event suggests its possible role in driving early tumorigenesis, with later acquisition of a subclonal *BRCA2* p.R2341C mutation (Fig. 4A). Both BAP1 and BRCA2 are implicated in DNA damage repair and inactivating mutations in these genes can contribute to sensitivity to DNA-damaging[32, 33].These observations provide a biological rationale for this patient’s exceptional response to Gem/Doce.

**Figure 4.**
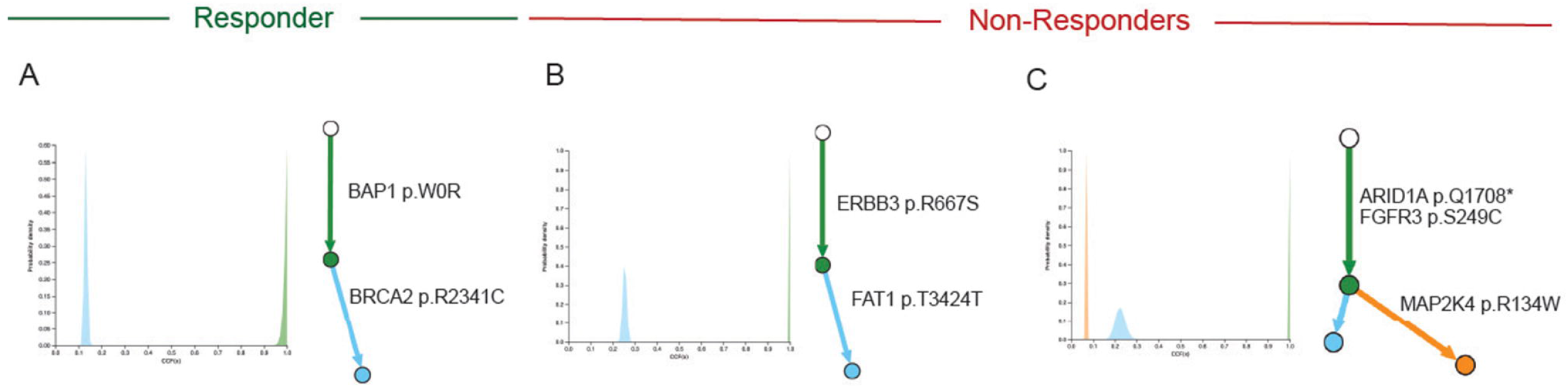
Genomic evolution in responder and non-responder tumors to Gem/Doce. Histograms with cancer cell fractions (CCFs) for clonal mutations are represented in green and subclonal mutaions in blue and orange. For each tumor, the corresponding phylogenetic tree is shown on the right of the histogram. Select cancer genes are annotated in the phylogenetic trees A. Tumor from a responder to Gem/Doce harbor mutations in genes involved in homologous recombination, specifically a loss-of-function truncating BAP1 p.W0R clonal mutation and a BRCA2 p.R2341C subclonal mutation. B. In a non-responder case, there is an oncogenic ERBB3 p.R667S clonal mutation and FAT1 p.T3424T subclonal mutation that leads to aberrant activation of the Wnt/ β-catenin pathway. C. In another non-responder tumor, there is loss-of-function truncating ARID1A p.Q1708 clonal mutation as well as oncogenic activating FGFR3 p.S249C clonal mutation.

In the two non-responder cases, the clonal trees were defined by both clonal and subclonal alterations not implicated in DNA repair. For example, in one non-responder patient tumor sample we observed a clonal p.R667S mutation in the kinase domain of *ERBB3* (HER3) (Fig. 4B). Interestingly, in the other non-responder case we observed a branched clonal tree and a clonal p.S249C mutation in the tyrosine kinase receptor *FGFR3* that is implicated in chemoresistance in bladder cancer cells (Fig. 4C)[34]. This case also had a clonal truncating mutation in *ARID1A* p.Q1708*, a subunit of the SWI/SNF chromatin remodeling complex. *ARID1A* inactivation is associated with progression and aggressiveness in bladder cancer[11, 35].

The presence of clonal and subclonal mutations in DNA damage repair genes further supports the hypothesis that tumors responsive to Gem/Doce are extremely sensitive to cytotoxic chemotherapy, resulting in prolonged high-grade recurrence-free and cystectomy-free survival in these patients (Fig. 1). In contrast, non-responders have mutations in tyrosine kinases (e.g. *FGFR3*) that may benefit from specific targeted therapies available for NMIBC.

## Discussion

Gemcitabine/Docetaxel has emerged as an effective regimen with favorable efficacy and tolerability after BCG failure, but no predictive biomarkers have been identified to guide patient selection. In this study, we used whole-exome sequencing to define the mutational features of BCG unresponsive NMIBC and nominate genomic drivers of response and resistance to Gem/Doce. To our knowledge, this is the first study to comprehensively integrate genomic alterations and response to Gem/Doce in patients with BCG unresponsive NMIBC. We found that responders harbored mutations in DNA damage repair gene such as *BRCA1* and *BRCA2* which might explain cancer cell sensitivity to intravesical Gem/Doce. DNA damage repair alterations in genes are increasingly recognized as important determinants of NMIBC tumor biology. For example, *BRCA2* mutations are associated with decreased recurrence and progression in patients with treatment naive high risk NMIBC[36]. However, the association between these mutations and response to treatment, specifically Gem/Doce, was not previously established. Moreover, our phylogenic analysis revealed clonal and subclonal mutations in *BAP1* and *BRCA2*, genes involved in homologous recombination, further supporting a potential biological role for defective DNA damage repair in mediating response to Gem/Doce. Lastly, our results identified STAG2 and CREBBP mutations exclusively in responders. Larger NMIBC cohorts identified mutation rates of 24% for STAG2 and 20% for CREBBP[13], suggesting biological roles for these genes in bladder cancer pathogenesis.

Although non-responders exhibited a significantly higher overall TMB, these tumors did not carry more alterations in DNA damage repair genes. Previous studies have suggested a potential association between TMB and progression to MIBC[36, 37], but the mean number of mutations in those studies is broad, reflecting the underlying heterogeneity of NMIBC and different sequencing platforms. Interestingly, in our cohort, non-responders had more frequent whole-genome duplication and APOBEC mutational signatures, which have been implicated in resistance to cytotoxic chemotherapy[38], harbored mutations associated with a worse prognosis in bladder cancer (e.g. *ERBB3* and *ARID1A*), and demonstrated more complex branched patterns of clonal evolution, underlying different mechanisms of resistance that can be potentially overcome with targeted therapies. For example, we identified clonal *FGFR3* actionable mutations (e.g. *FGFR3* p.S249C) targeted by the FGFR inhibitor erdafitinib[39]. These results highlight the importance of molecular profiling of BCG-unresponsive NMIBC prior to second line intravesical therapies because some mutations present in non-responders to Gem/Doce (e.g. *FGFR3*) can be specifically targeted with other therapies.

Our study has limitations. First, our sample size was modest, limited by the small amounts of bladder tissue obtained at time of TURBT in line with challenges experienced by several groups[40]. Additionally, germline DNA for matched tumor-normal analysis was not available in several patients. However, three patients had matched germline DNA and genomic features as tumor mutation burden and single nucleotide variant cells were similar with tumor only samples, which indicates a harmonized analysis of two approaches.

## Conclusions

Together, our results indicate the genomic landscape — particularly DNA repair capacity and clonal architecture — shapes BCG unresponsive tumors’ vulnerability to Gem/Doce. Future studies with larger multi-institutional cohorts will be essential to validate DDR gene alterations as predictive biomarkers of response to Gem/Doce and prospectively test targeted therapeutic combinations (e.g. FGFR inhibitors) for patients with resistant disease. Integration of genomic profiling into clinical decision-making may ultimately tailor selection of intravesical therapies and improve outcomes for patients with BCG-unresponsive NMIBC.

## Take Home Message

Whole-exome sequencing of BCG-unresponsive NMIBC identified genomic differences underlying response to Gem/Doce. Tumors with DNA damage repair alterations are extremely sensitive to Gem/Doce and patients experience durable recurrence-free survival. In contrast, BCG-unresponsive tumors resistant to Gem/Doce have genomic instability and targetable mutations (e.g. FGFR3) that support biomarker-driven treatment strategies in the future.

## Author statement

**Kendrick Yim:** Conceptualization, Methodology, Software, Formal analysis, Investigation, Data curation, Visualization, Writing-Original draft preparation, Reviewing and Editing. **Matias Vergara:** Conceptualization, Methodology, Software, Formal analysis, Investigation, Data curation, Visualization, Writing-Original draft preparation, Reviewing and Editing. **Jihyun Lee:** Formal analysis, Investigation, Writing-Reviewing and Editing. **Brendan Reardon**: Formal analysis, Investigation, Writing-Reviewing and Editing. **Jihye Park:** Formal analysis, Investigation, Writing-Reviewing and Editing. **Kevin Melnick:** Data curation, Writing-Reviewing and Editing. **Timothy N Clinton**: Data curation, Writing-Reviewing and Editing. **Matthew Mossanen:** Data curation, Writing-Reviewing and Editing. **Graeme S Steele**: Data curation, Writing-Reviewing and Editing. **Jessica Bolduc**: Data curation, Writing-Reviewing and Editing. **Michelle Hirsch:** Data curation, Writing-Reviewing and Editing. **Natalie Rizzo:** Data curation, Writing-Reviewing and Editing. **Chin-Lee Wu**: Data curation, Writing-Reviewing and Editing. **Matthew Wszolek**: Data curation, Writing-Reviewing and Editing. Keyan Salari: Data curation, Writing-Reviewing and Editing. **Adam Feldman**: Data curation, Writing-Reviewing and Editing. **Adam S Kibel**: Data curation, Writing-Reviewing and Editing. **Kent W Mouw**: Conceptualization, Methodology, Investigation, Writing-Original draft preparation, Reviewing and Editing, Funding acquisition. **Eliezer Van Allen:** Conceptualization, Methodology, Investigation, Writing-Original draft preparation, Reviewing and Editing, Funding acquisition. **Mark A Preston:** Conceptualization, Methodology, Investigation, Writing-Original draft preparation, Reviewing and Editing, Funding acquisition. **Filipe LF Carvalho:** Conceptualization, Methodology, Investigation, Writing-Original draft preparation, Reviewing and Editing, Supervision, Project administration, Funding acquisition.

## Acknowledgments

We thank the patients who participated in this study and their families. This work was supported by the National Cancer Institute (R01 CA279221 (E.M.V.A., K.W.M.), R01CA272657 (K.W.M.), P30 CA006516 (F.L.F.C.), K08 CA282969 (F.L.F.C.)), Parker Institute for Cancer Immunotherapy (E.M.V.A.), Bladder Cancer Advocacy Network (F.L.F.C.) and Karin Grunebaum Cancer Research Foundation (F.L.F.C.).

