## Supplementary figures and images for "Genomic analysis of BCG unresponsive non-muscle-invasive bladder cancer identifies drivers of sensitivity to intravesical Gemcitabine/Docetaxel"

### Supplemental Figure 1

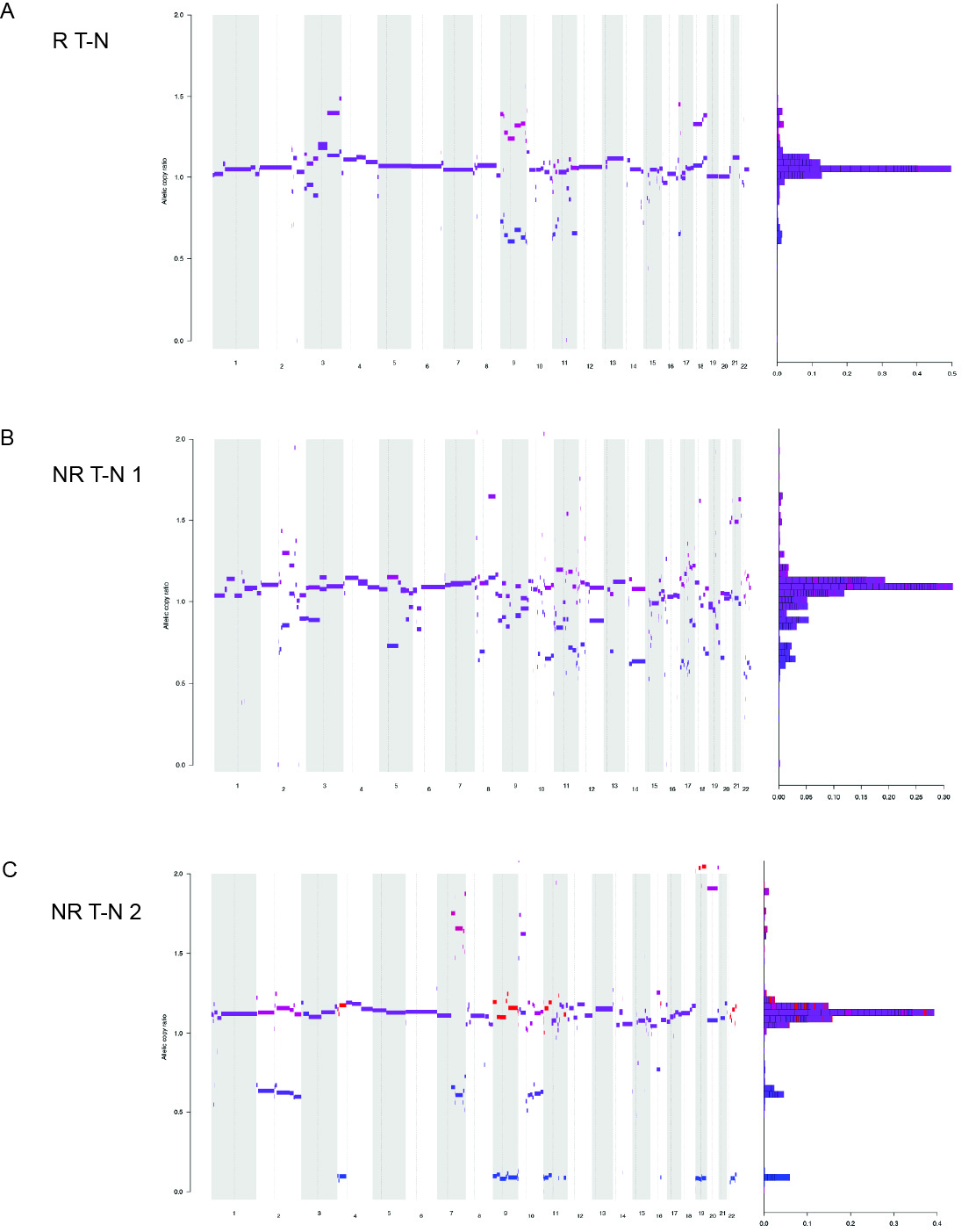

### Supplemental Figure 2

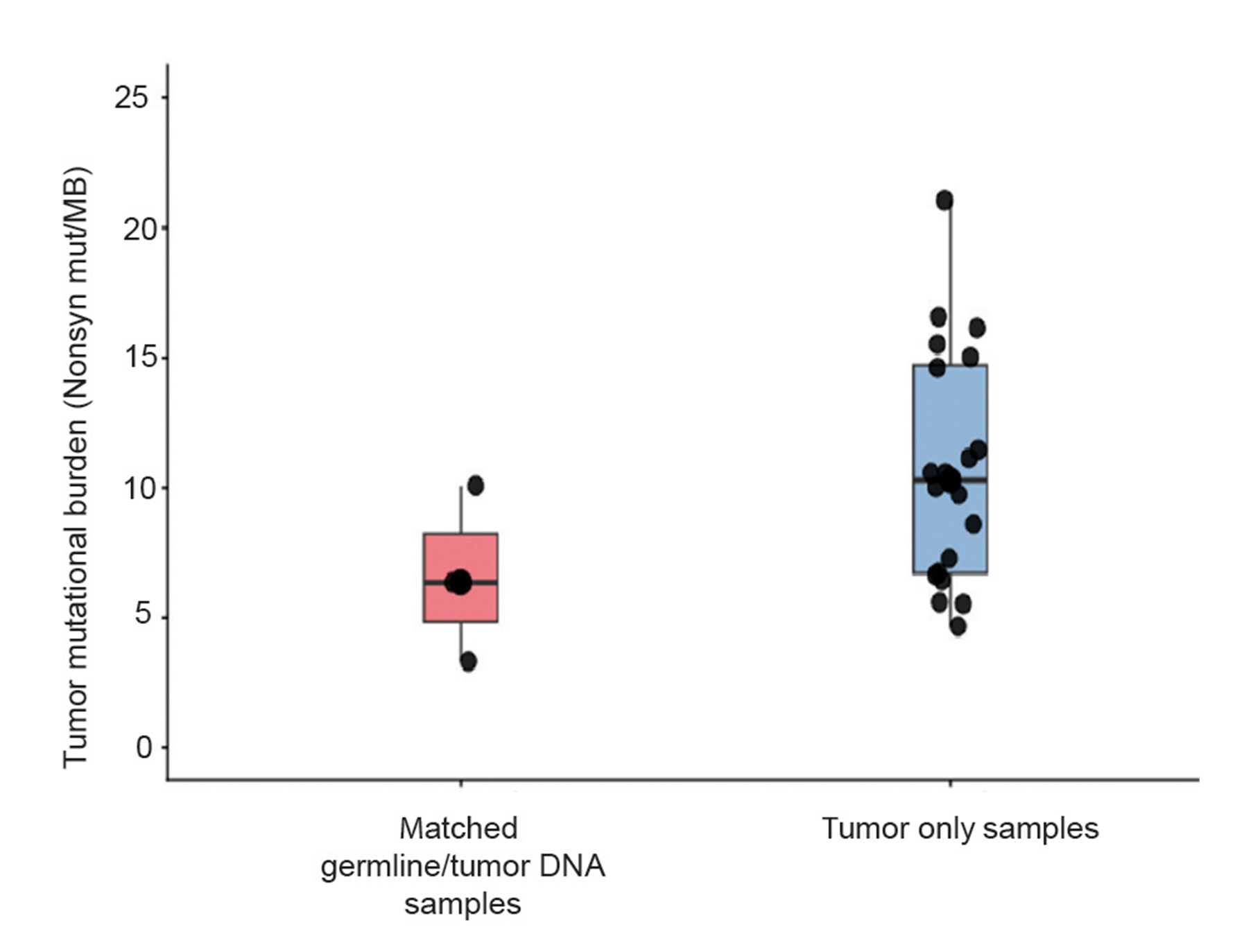

### Supplemental Figure 3

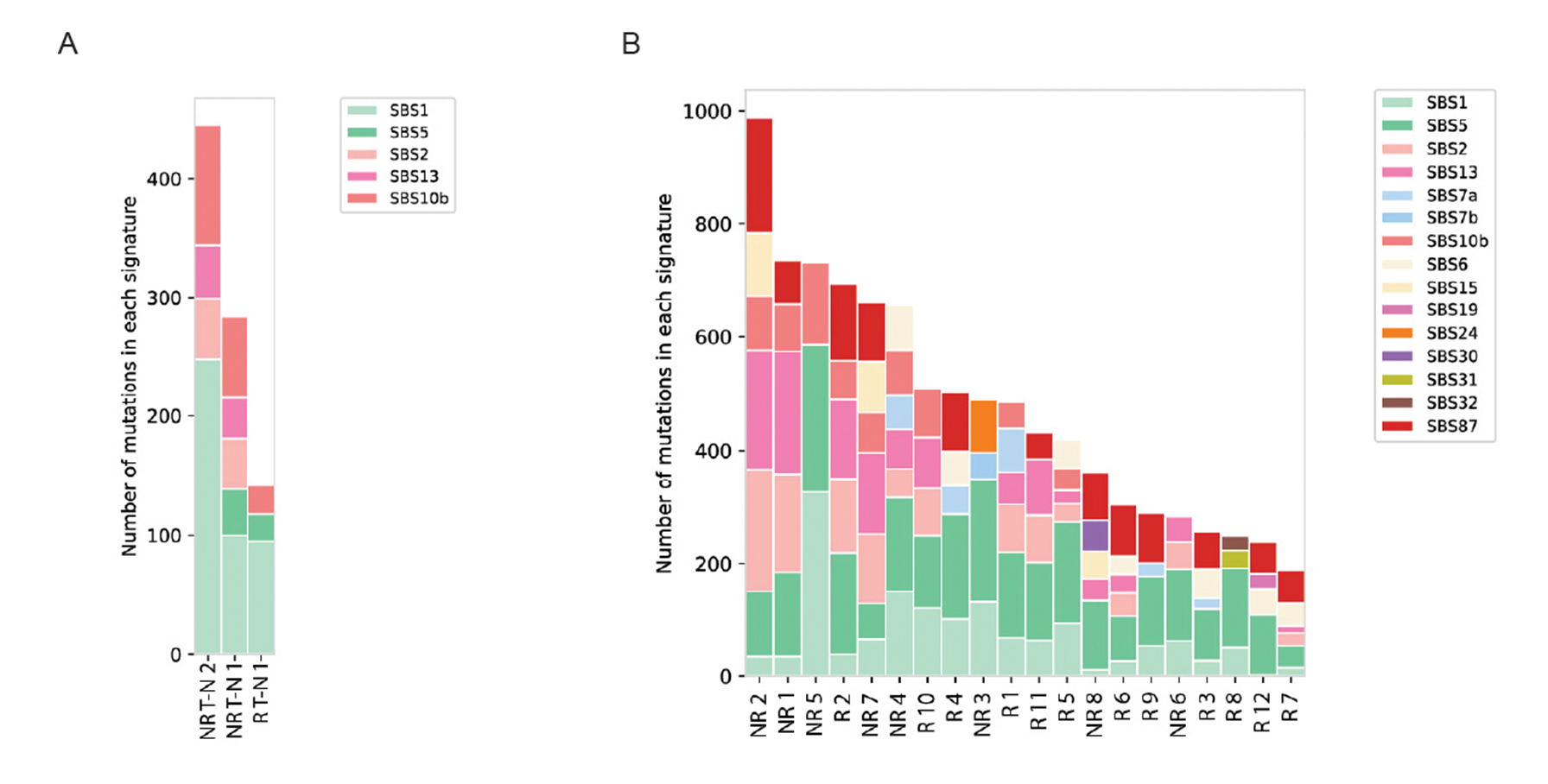
